# A dNmnat-sensitized *in vivo* platform for unbiased discovery of regulators of neurodegeneration

**DOI:** 10.1101/2025.09.25.678663

**Authors:** Adel Avetisyan, Ernesto Manzo, Romina Barria, Magdalena Kocia, Jiun-Min Hsu, Alexandra Larson, Rachel Weirnick, Kavya Balasubramanian, Nathan Blackwell, Micheal Freeman, Dana Marie Bis, Sarah Fazal, Stefan Schüchner, Lukas Neukomm, Marc Freeman

## Abstract

Axon degeneration drives nervous system dysfunction in diverse neurological diseases and injuries. While key regulators of injury-induced Wallerian degeneration have been identified, approaches have largely relied on axotomy, a particularly extreme injury for a neuron. Here, we developed a forward genetic screening platform in the adult *Drosophila* wing that sensitizes neurons to degeneration through depletion of nicotinamide mononucleotide adenylyltransferase (dNmnat), the essential NAD^+^ biosynthetic enzyme. We screened 9,393 mutagenized chromosomes and recovered 59 mutations that suppress neurodegeneration induced by dNmant depletion, including the known pro-degenerative molecule dSarm, thereby validating our approach. We show loss of *CG4098*, the *Drosophila* homolog of the mammalian enzyme NUDT9, robustly preserved axons and induced widespread remodeling of NAD^+^-related metabolites, which provides a new link between cADP-ribose hydrolase activity and neuronal survival. We further show mutations in the transcription factor Abrupt resulted in elevated dNmnat protein levels and blocked axon degeneration, suggesting Abrupt normally tunes dNmnat levels and neuronal NAD^+^ homeostasis. dNmnat depletion is thus a versatile approach for unbiased discovery of axon death pathway components, each of which may provide new entry points for therapeutic strategies to preserve axons in neurodegenerative diseases.

## Introduction

Axon degeneration is a hallmark of many neurological disorders, including multiple sclerosis^1^, spinal muscular atrophy^2^, amyotrophic lateral sclerosis^3^, and Parkinson’s disease^4^. Axon loss also occurs in response to toxins^5,6^, chemotherapy^7^, diabetes^8^, and myelin disorders^9^. Loss of axons irreversibly disrupts neuronal circuits and compromises nervous system function. Identifying the molecular pathways that drive axon degeneration is essential for developing strategies to preserve axons in disease and injury.

Genetic studies in *Drosophila* have been instrumental in identifying conserved regulators injury-induced of axon degeneration, such as *dSarm, highwire* and *axundead*^10–15^. Loss of function alleles of *dSarm* were discovered to potently suppress axon degeneration in flies, and subsequently mouse (*SARM1*), after axotomy^12^, where axons are physically severed from their cell bodies. While powerful, highly reproducible and easy to execute, axotomy is among the most severe of axonal injuries. When used as a tool to study axon degeneration, this approach may enable detailed analysis of only the most potent suppressors of axon degeneration –mutations that are capable of keeping axons intact for extended periods of time after loss of all support from their cell bodies. Physiologically relevant regulators of axon degeneration that act in other contexts that might be less severe in nature (e.g. dying back neuropathies) may therefore go undetected. Moreover, axon degeneration in chronic neurodegenerative diseases occurs for the most part when axons remain attached to their cell bodies. *SARM1* mutations have indeed been shown to suppress axon loss in preclinical models of traumatic axonal injury, CIPN, DIPN, toxin exposure and a number of other preclinical models of neurodegenerative disease^16–21^. However, the extent to which axotomy serves as a broadly predictive model for axon loss in other contexts remains an open question.

Loss of nicotinamide mononucleotide adenylyltransferase (Nmnat), the rate-limiting enzyme in NAD^+^ synthesis, provides an alternative, genetically tractable trigger for neurodegeneration. Genetic depletion of the cyto- and axoplasmic Nmnat2 *in vitro* results in spontaneous axon degeneration in the absence of injury, and mice lacking *Nmnat2* exhibit perinatal death and short axons^22–24^. Remarkably, these phenotypes are completely suppressed by *SARM1* null mutations^25^; and genetic depletion of dNmnat in *Drosophila* neurons leads to spontaneous neurodegeneration that can be suppressed by *dSarm* null mutations^13–15,23,24^. These observations reveal that loss of dNmnat/Nmnat2 in neurons is sufficient to activate the dSarm/SARM1-dependent axon degeneration pathway, even in the absence of axotomy. Notably, degeneration caused by axotomy or dNmnat loss cannot be suppressed by inhibiting apoptosis, autophagy, or many other pro-degenerative pathways, suggesting the dSarm/SARM1 pro-degenerative pathway is genetically distinct ^12,13,26,27^.

Here we report our development of a high-throughput, unbiased *in vivo* forward genetic screening approach in the adult *Drosophila* wing to identify genes required to drive axon degeneration. Our method enables imaging neurodegeneration with single cell resolution in live animals, examination of degeneration in axons, cell bodies and dendrites, rapid forward genetic screening without tissue dissection. We report the results of a screen of 9,393 chromosomes, which led to the identification of 59 mutants that suppress neurodegeneration after dNmnat depletion. We mapped one suppressor CG4098 (NudT9) to the NAD^+^ metabolic pathway, and identify the transcription factor Abrupt as a regulator of dNmnat levels. While both strongly suppresses neurodegeneration after dNmnat loss, neither suppress axon degeneration after axotomy. We argue this new approach has the potential to uncover a broader and perhaps more physiologically relevant set of axon death regulators, and ultimately lead to new strategies to block axon degeneration in disease and injury.

## Results

### Activation of dSarm by dNmnat depletion

We previously showed that neuronal depletion of dNmnat by RNAi led to neurodegeneration that could be suppressed by null alleles of *dSarm* or *axundead*. To identify genetic suppressors of dSarm-dependent axon degeneration across the *Drosophila* genome, we developed a forward genetic screening platform using the adult fly wing (Fig. 1A). In this system, we activated neurodegeneration by targeted knockdown of *dNmnat* via RNA interference (RNAi) in individual peripheral sensory neurons of the *Drosophila* wing. The L1 sensory nerve in the adult wing houses ∼280-290 peripheral sensory neurons^28^. Neuron-specific expression was achieved using the Gal4/UAS system in combination with MARCM (Mosaic Analysis with Repressible Cell Marker) technique^29^. Neuronal clones were generated using a neuronal specific Gal4 driver in combination with the *ase-Flp*^13^ recombinase, which resulted in the production of small numbers of neuronal clones expressing GFP and *dNmant*^*RNAi*^ in the adult wing.

**Figure 1.**
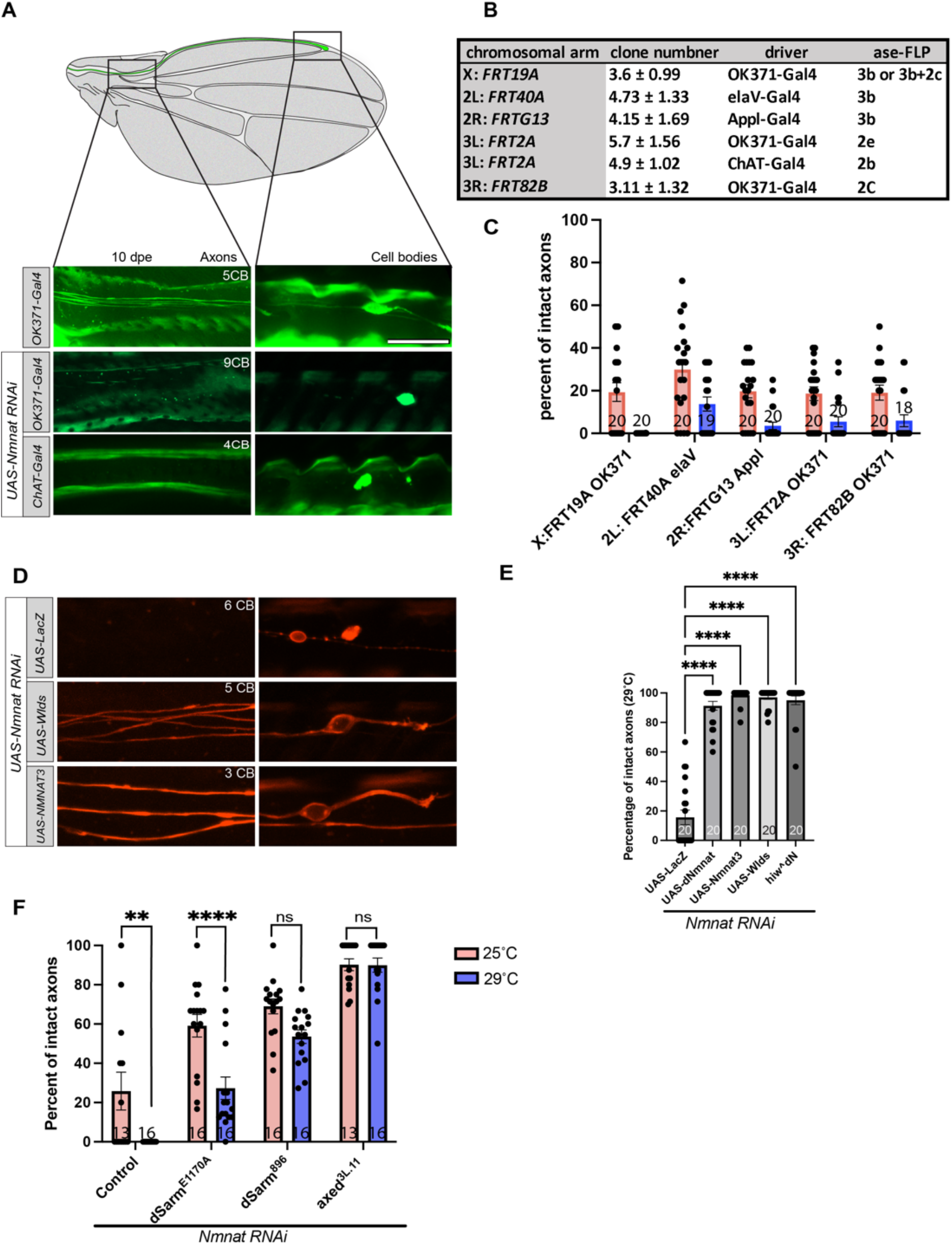
dNmnat depletion triggers axon degeneration in the adult Drosophila wing. (A) Schematic of the adult wing showing the L1 sensory nerve region analyzed. Representative confocal images of neuronal clones at 10 days post-eclosion (dpe) illustrate robust axon degeneration upon expression of dNmnat RNAi (*OK371>UAS-dNmnat RNAi*) compared with controls. Axonal and cell body regions are shown. (B) MARCM-based strategy enables generation of neuronal clones on all major chromosomal arms. Table summarizes clone number (mean ± s.e.m.), Gal4 drivers, and *ase-FLP* insertion used. (C) Quantification of intact axonal zones in *dNmnat*^*RNAi*^ expressing clones at 25 °C and 29 °C, showing enhanced degeneration at elevated temperature (*p < 0.05, **p < 0.01, ***p < 0.001; Student t-test; sample size denoted below bar graph). (D) Representative images of wings expressing *dNmnat*^*RNAi*^ alone or in combination with *UAS-Nmnat3* or *UAS-Wld*^*S*^. NMNAT3, and Wld^S^ overexpression restores axon integrity. *LacZ* is used as a control to account for the number of UAS constructs. (E) Quantification of intact axons by dNmnat, Nmnat3, Wld^S^, or hiw^dN^ in neuronal clones. (F) Suppression of axon degeneration by *Sarm*^*E1170A*^, *dSarm*^*896*^, and *axed*^*3L*.*11*^ quantified.

In general, we aimed to induce axon degeneration by ∼10 days post-eclosion (dpe) (Figure 1A). However, as in our previous work, we found variability in clone production and rates of neurodegeneration on different chromosomes, and with different *Gal4* driver lines^13,14,28,30^. We therefore systematically characterized combinations of Gal4 drivers and FRT sites with the goal of identifying combinations for each arm of each major chromosome that (1) produce a small number of clones (between 2-9 clones) and (2) potently induce axon degeneration by 10 dpe (Fig. 1A, 1B, 1C). We were able to assemble lines to target the X chromosome, 3L and 3R using the glutamatergic *OK371-Gal4* or cholinergic *ChAT-Gal4* driver (Fig. 1B); and 2L and 2R using the panneuronal drivers *elaV-Gal4* and *Appl-Gal4* (Fig. 1B); thereby gaining access to the entire *Drosophila* genome with the exception of chromosome 4 and regions of each chromosome near centromeres that are proximal to FRT sites (see methods).

dNmnat knockdown by *UAS-dNmnat*^*RNAi*^ induced dying-back neurodegeneration, characterized by the propagation of degeneration from synaptic terminals and distal axons toward the cell bodies (Supp Figure X) and resulted in markedly reduced axon survival (mean survival rate of: 15.6%; Fig. 1D, 1E). Because the Gal4/UAS system is temperature sensitive^31^, we hoped to be able to increase the efficiency of the *UAS-dNmnat*^*RNAi*^, and thereby neurodegeneration, by increasing the temperature flies were reared at to 29^°^C^32^. Indeed, we found rearing animals at 29^°^C resulted in a significant decrease in the number of intact clones after 10 dpe for all lines tested (Fig. 1C). We next examined whether replenishing dNmnat activity in neuronal clones could protect them from degeneration by expressing *dNmnat*, mouse *NMNAT3*, or *Wld*^*S*^, which we previously showed robustly protect axons from degeneration following axotomy^33^. Expression of each of these molecules resulted in robust protection of *dNmnat*^*RNAi*^ clones (mean survival rates: 91.28%, 98.44%, 97.04%, respectively; Fig. 1D and 1E). We and others have previously reported that loss of function mutations in *highwire* (*hiw*) were sufficient to protect axons from degeneration^10,14^. Highwire is believed to drive UPS-mediated degradation of dNmnat, with loss of Highwire leading to stabilization of dNmnat and protection of axons. While we predicted that loss of Highwire would not suppress dNmnat^RNAi^, we found that a dominant negative version of Highwire (Hiw^DN^) potently suppressed *dNmnat*^*RNAi*^ (mean survival rate: 95%; Fig 1D,E). These data suggest that dNmnat depletion, even after *dNmant*^*RNAi*^, is mediated by Highwire and does not require axotomy. Finally, to confirm that our knockdowns of dNmnat activated dSarm-dependent Wallerian-like degeneration, we confirmed that loss of function mutations in *dSarm*^12^ and *axundead*^14^ provided significant protection of both axons and cell bodies after dNmnat depletion, even at 29 °C (Fig. 1F).

Together these results demonstrate we can selectively target *dNmnat* knockdown in MARCM clones to activate dSarm-mediated axon degeneration *in vivo*, and that this system provides genetically tractable assay for identifying additional molecules that play a role in dSarm-mediated axon degeneration.

### Unbiased F_1_ forward genetic screening platform

Using the above genetic tools we performed an F_1_-based genetic mosaic screen to identify genetic modifiers of axon degeneration after dNmnat depletion in MARCM clones. P_0_ males were mutagenized with ethyl methanesulfonate (EMS) and subsequently crossed to virgin “tester” females carrying the appropriate MARCM genetic background (Fig. 2A). Within GFP^+^ clones we examined in the wing, cells were homozygous mutant for random EMS-induced mutations, while most other cells in the animal remained heterozygous. This allowed us to screen for modification of the *dNmnat*^*RNAi*^ phenotype in clones, even when mutations might otherwise cause animal lethality when homozygous throughout the animal.

**Figure 2.**
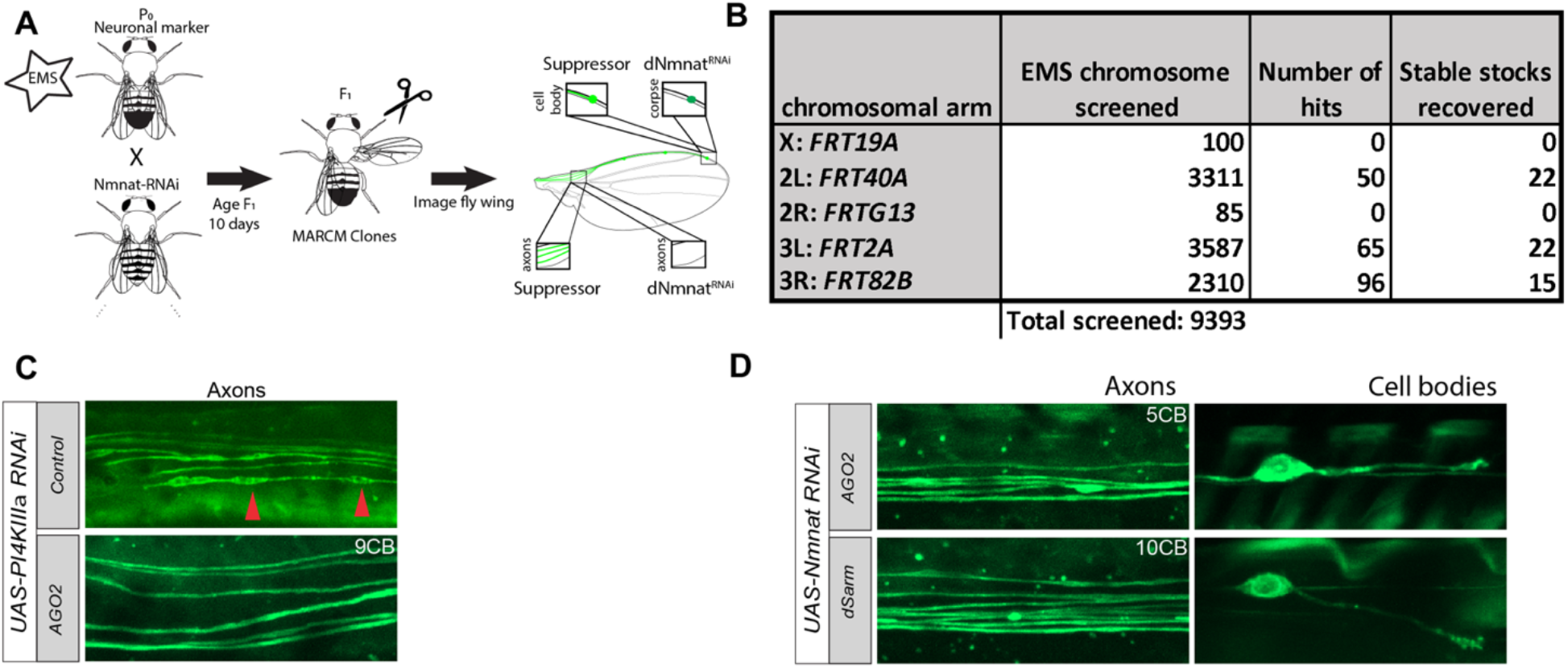
F1 forward genetic screen identifies suppressors of dNmnat-induced axon degeneration. (A) Schematic of the screening strategy. EMS-mutagenized P_0_ males were crossed to tester females carrying MARCM insertions and *UAS-dNmnat RNAi*. F_1_ progeny were aged for 10 days post-eclosion (dpe), and axons within the wing nerve were imaged to identify suppressor clones that maintained axonal integrity compared with wild-type controls. (B) Summary of EMS mutagenesis screen. A total of 9,393 chromosomes were screened across all major chromosomal arms. The number of hits and stable stocks recovered are indicated. (C) Testing the efficiency of the RNAi machinery. Representative confocal image of a *UAS-PI4KIIIαRNAi* clone shows robust axonal blebbing (red arrowheads), a phenotype used to distinguish true neuroprotective suppressors from false positives. Whole-genome sequencing of recovered lines identified *Ago2* as a false-positive hit due to impaired RNAi function. (D) Validation of candidate suppressors. Whole-genome sequencing of recovered lines identified *dSarm* as a strong suppressor.

F_1_ progeny were collected on the day of eclosion, aged for 10 days at either 25 °C or 29 °C, and then one wing was removed to assess the phenotype by light microscopy. We screened for mutations that resulted in preservation of distal axons and cell bodies in males (Fig. 2A) (i.e. to avoid having to collect virgin females). When suppression of neurodegeneration at day 10 was observed, the corresponding male was crossed to virgin females to establish a stable stock. The second wing of these males was examined five days later (at 15 dpe), to determine whether neuroprotection was still present. Some males exhibited sterility, a side effect of EMS treatment; however, despite these challenges, 15–44% of the originally screened mutant chromosomes that exhibited suppression of neurodegeneration in *dNmant*^*RNAi*^ backgrounds were successfully recovered. In total, we screened 9,393 mutagenized chromosomes across all major chromosomal arms of the *Drosophila* genome and recovered 59 lines for further analysis (Fig. 2B). Although we retained all of the strongest suppressors of neurodegeneration, only two mutants that were new loss of function alleles of *dSarm* (see below), were able to protect severed axons from Wallerian degeneration caused by axotomy. All other suppressors failed to block Wallerian degeneration.

It is possible that we might identify loss of function mutations in the RNAi machinery, which would result in suppression of neurodegeneration phenotypes associated with *dNmnat*^*RNAi*^. To distinguish true neuroprotective backgrounds from artifacts caused by impaired RNAi activity, we examined whether mutant clones retained the ability to form axonal blebs that occur after PI4KIIIα knockdown. In control neuronal clones, and most suppressors, PI4KIIIα RNAi consistently induced robust axonal blebbing, indicating that RNAi activity remained efficacious. However, we identified two mutants that failed to results in blebbing of axons (Fig. 2C). Whole genome sequencing (WGS) demonstrated these lines have mutations that caused a premature stop codon in *Ago2*: line *3L*.*1696 –* Q29* (Fig. 2C) and *line 3L*.*2494 –* Q999*. Since *Ago2* is a core component of the RNAi machinery in *Drosophila*, the *Ago2* multination suggests that alterations to the RNAi machinery can give rise to false positives.

All additional suppressors of *dNmnat*^*RNAi*^-induced neurodegeneration were subjected to WGS to identify potential mutations responsible for axon protective phenotypes. Two hits— those that also protected axons from axotomy-induced axon degeneration—harbored missense mutations in *dSarm*: one within the genomic region corresponding to the ARM domain, resulting in substitution of the evolutionarily conserved P924 with leucine (numbering according to dSarm-PI), which corresponds to P298 in mouse and human SARM1; and another within the TIR domain, causing substitution of the conserved V1216 with methionine, corresponding to V635 in mouse and V595 in human SARM1, respectively.

These findings demonstrate that our screen can identify strong suppressors of axon degeneration, including new alleles of *dSarm*. While the vast majority suppressed *dNmant*^*RNAi*^-induced neurodegeneration, none (except *dSarm* mutants) were capable of suppressing axon degeneration after axotomy. This argues that while neurodegeneration after loss of dNmnat activity or axotomy share many features and genetic components (e.g. *dSarm, axundead, hiw*), they are genetically distinguishable events. Below we describe our more detailed molecular analysis of two new mutants.

### Mutations in the ADP-ribose hydrolase CG4098 protect axons from *dNmant*^*RNAi*^-induced neurodegeneration

In the above screen we identified an EMS mutant line (*3L0184*) that markedly increases the survival of *dNmnat*^*RNAi*^-expressing neuronal clones from 18.61% to 60.58% (Fig. 3A). Whole-genome sequencing of *3L0184* revealed a 57-base pair deletion predicted to cause a frameshift and premature stop codon insertion of the uncharacterized *Drosophila* gene *CG4098*. CG4098 is the homolog of human NUDT9 (nudix hydrolase 9, DIOPT v9.1 score: 14/14), a ubiquitously expressed enzyme that hydrolyzes ADP-ribose (ADPR) to AMP and ribose-5′-phosphate^34^. ADPR is a metabolite of NAD^+^ consuming enzymes, e.g. dSarm/SARM1, and elevated AMP levels have been implicated in mitochondrial dysfunction^35^. Given the established role of NAD^+^ in the axon degeneration pathway and the potential contribution of mitochondrial dysfunction to neurodegenerative disease^36^, *CG4098* emerges as a compelling candidate for further functional characterization. To confirm we identified the correct gene, we generated a CRISPR/Cas9 induced frameshift allele of *CG4098* containing a 358 base pair deletion and an adenine insertion (referred as *CG4098*^*KO*^; Supplementary Fig.1). *CG4098*^*KO*^ clones were also strongly protected from dNmnat depletion (mean axon survival: 53.7%; Fig. 3A).

**Figure 3.**
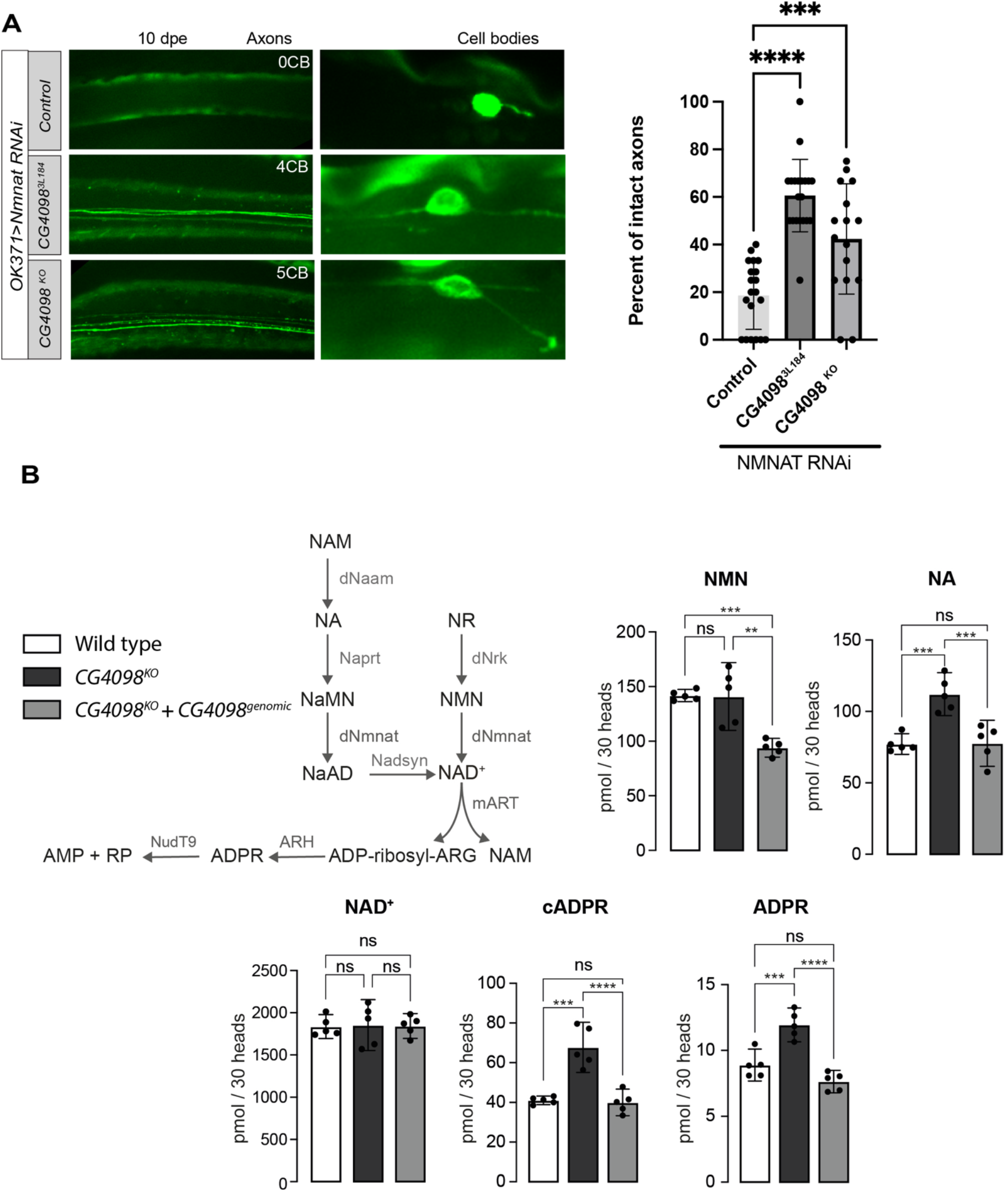
Loss of CG4098/NUDT9 protects axons from degeneration and alters NAD^+^-related metabolites. (A) Representative confocal images of adult wing axons and cell bodies at 10 days post-eclosion (dpe) from control, *OK371>dNmnat RNAi*, EMS mutant line *CG4098*^*3L184*^, and *CG4098* knockout (*CG4098*^*KO*^) clones. Quantification (right) shows significant protection of axons in *CG4098*^*3L184*^ and *CG4098*^*KO*^ backgrounds compared with controls (***p < 0.001, ****p < 0.0001; one-way ANOVA). Scale bar, 20 μm. (B) Schematic of NAD^+^ metabolic pathways and metabolites linked to CG4098/NUDT9 function (left). Metabolite profiling of adult fly heads from wild-type, *CG4098*^*KO*^, and *CG4098*^*KO*^ + *CG4098*^*KO*^+ *CG4098*^genomic^ reveals significant changes in cADPR and ADPR, but not NAD^+^ or NA. Quantification (right) shows increased NMN and ADPR, and accumulation of cADPR in *CG4098*^*KO*^ mutants, which are restored in rescue conditions. Data are mean ± s.e.m.; ***p < 0.001, ****p < 0.0001; ns, not significant.

Given the importance of NAD^+^ metabolism in axon degeneration, and the fact that dSarm/Sarm1 is an NAD^+^ hydrolase, we reasoned that alterations in the NAD^+^ metabolism may contribute to the protective nature of *CG4098*^*KO*^ clones after dNmnat depletion. We therefore profiled metabolites in adult *Drosophila* heads comparing (1) the w^1118^ control background line, (2) *CG4098*^*KO*^, and (3) *CG4098*^*KO*^ with a genetic BAC clone that contains the CG4098 gene as a rescue (Fig. 3B, all three genotypes without dNmnat depletion). While total NAD^+^ levels are not changed, the substrates consumed by NudT9/CG4098 - cADPR and ADPR – were elevated in *CG4098*^*KO*^ conditions and restored in the genetic rescue condition (Fig. 3B). These results support the notion that flies lacking *CG4098* do not consume cADPR and ADPR as efficiently as controls. Similarly, Nicotinic acid (NA) and Nicotinic acid riboside (NaR) were elevated in *CG4098*^*KO*^ lines and restored in the rescue line (Fig. 3C and Supplementary Fig. 2). Changes in NMN were also significantly decreased in the genetic rescue condition (Fig. 3B). Together, these data indicate broad remodeling of metabolites linked to NAD^+^ synthesis and turnover in the absence of *CG4098* (Supplementary Fig. 2).

### Loss of function alleles of *abrupt* protects neurons from neurodegeneration induced by dNmnat depletion

We also identified line *2L153*, which significantly protected neuronal clones from neurodegeneration after dNmnat depletion (Fig. 4A). WGS of line 2L153 identified a G>A mutation at location 2L:11211964, which results in a splice donor variant between exons 2 and 3. The *abrupt* gene encodes a BTB-zinc finger transcription factor previously found to be important in neuronal differentiation of dendritic arborization neurons^37^. To confirm this was the causative mutations, we tested an independent *abrupt* mutation, *ab*^*K02807*^, which contains a P-element insertion in the second intron of the gene and has been previously used as a loss of function allele^38^. We found that *ab*^*K02807*^ mutant clones potently protected axons from neurodegeneration in *dNmnat*^*RNAi*^ backgrounds (Fig. 4A).

**Figure 4.**
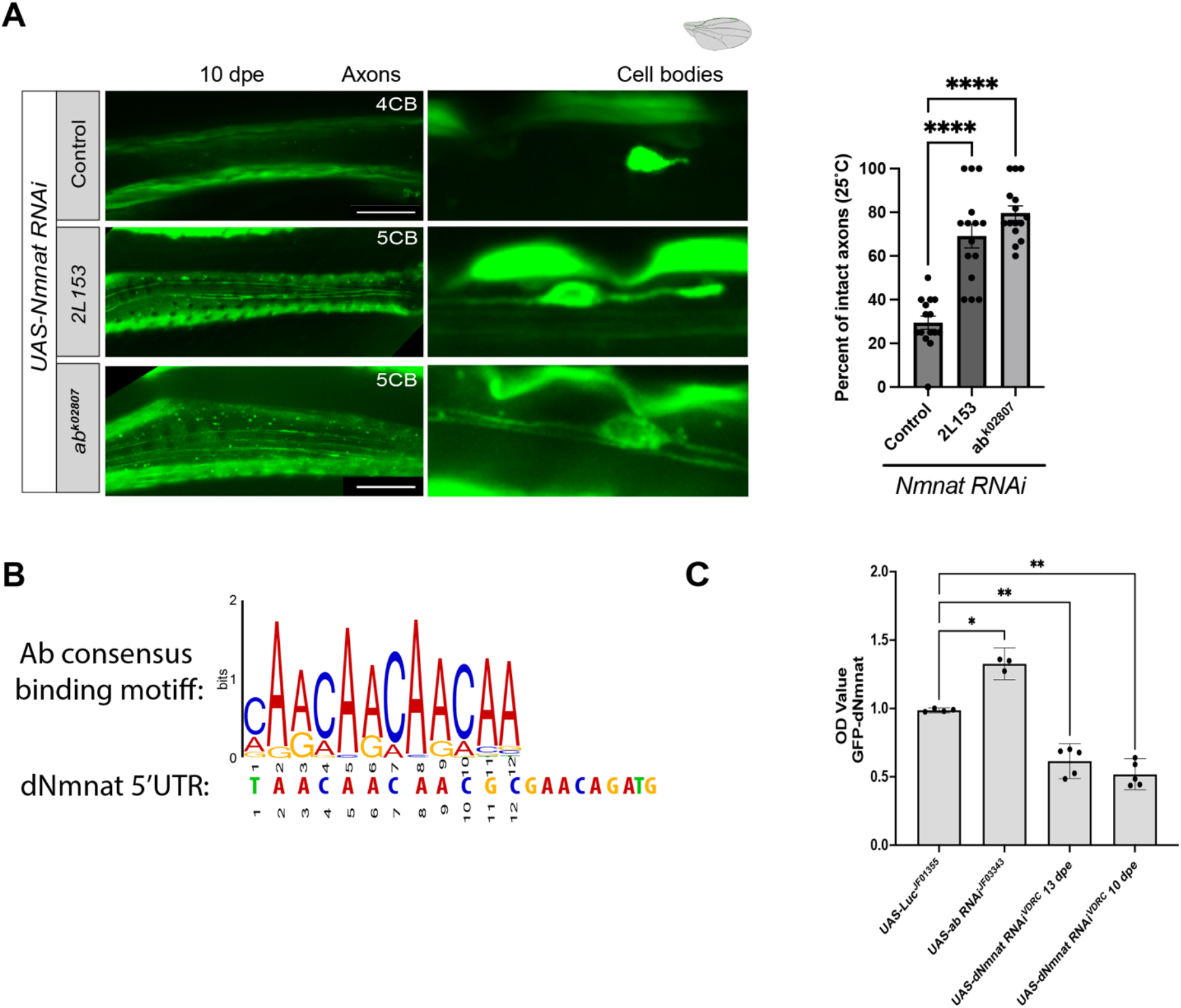
Loss of Abrupt protects axons from degeneration by regulating dNmnat expression. (A) Representative confocal images of adult wing axons and cell bodies at 10 days post-eclosion (dpe) from control, *OK371>dNmnat RNAi*, and Abrupt mutant clones (*ab*^*2L153*^ and *ab*^*K02807*^). Quantification (right) shows significant protection of axons in Abrupt mutant backgrounds compared with controls (****p < 0.0001; one-way ANOVA). Scale bar, 20 μm. (B) Sequence alignment of the Abrupt (Ab) consensus binding motif identified by ^39^ with the 5^ʹ^ untranslated region (UTR) of *dNmnat*, showing overlap with a predicted Ab binding site. (C) ELISA quantification of dNmnat-GFP protein levels in indicated genotypes. Knockdown of Abrupt significantly increases dNmnat levels compared with controls (*p < 0.05, **p < 0.01; one-way ANOVA). Data are mean ± s.e.m.

The *Drosophila* Abrupt consensus DNA binding site has been defined *in vivo*^39^, and, consistent with a potential role for Abrupt in regulation dNmnat directly, we identified a consensus binding motif in the 5’ UTR of *dNmnat* (Fig. 4B). Ab is best known as a transcriptional repressor^40^, suggesting that Ab might negatively regulate *dNmnat* through its binding site in the 5’ UTR. If true, we reasoned that the absence of Ab would cause dNmnat levels to increase. To test this hypothesis, we took advantage of the endogenously GFP-tagged dNmnat line^41^ and assayed whether knockdown of Ab would modify dNmnat-GFP levels in this background using ELISA assays^42^. Interestingly, we found that pan-neuronal knockdown of Ab significantly increased the levels of the GFP-tagged dNmnat compared to control (Fig. 4C), consistent with the notion that Ab mutants protect axons from degenerating through increased dNmnat levels.

In summary, our new approach successfully identified 59 new mutants that suppressed neurodegeneration after *dNmnat*^*RNAi*^. These included two new alleles of *dSarm*, validating the idea that we are targeting the dSarm pathway, a component of the NAD^+^ metabolic pathway (CG4098) and a potential transcriptional regulator of dNmnat. We propose that additional high throughput screening and subsequent thorough genetic analysis should lead to the identification of many new components of the dSarm/SARM1 pro-degenerative signaling pathway.

## Discussion

After injury, axons are capable of initiating an active molecular death pathway to drive degeneration. Over the past two decades, our understanding of how this process is initiated has deepened significantly. Genetic studies have established that loss of Nmnat enzymes initiate a self-destruction program mediated by dSarm/SARM1, and blocking this pathway can potently suppress degeneration^12–14,23–25^. However, most screens that have led to the identification of pro-degenerative molecules have relied on axotomy. While highly reproducible, axotomy is a severe injury that may not fully utilize the panoply of signals that can initiate axon degeneration *in vivo*, and screens using axotomy may therefore only reveal the strongest suppressors, which by definition must be capable of sustaining a distal axon stump in the absence of a cell body. Here, we developed a forward genetic screening platform using dNmnat depletion—a key activator of dSarm/SARM1-dependent neurodegeneration across many species—in the adult *Drosophila* wing to identify neuroprotective mutations. This approach uncovered known regulators of axon degeneration (Fig. 2D) and 57 potentially new neuroprotective mutations, including the novel modifiers of axon survival *CG4098* and *abrupt* (Fig. 3A; Fig 4A).

That loss of any single gene (e.g. *dSarm*/*Sarm1*) can lead to the long-term survival of distal axons after they are severed from the cell body is quite remarkable. This was first observed in the *Wld*^*S*^ mutant background, where a stabilized version of the NAD+ biosynthetic enzyme Nmnat1 is expressed^22,27,43,44^. However, it was reasonable to assume that overexpression of this molecule was neomorphic in nature, abnormally enabling maintenance of NAD^+^ and thereby survival. How an axon, once severed from its cell body, can maintain its integrity and functional properties for weeks in any mutant background remains unclear^12,13^. How many dSarm-like genes are there in the genome? We previously performed a forward genetic screens for mutants that would phenocopy *dSarm* nulls in axotomy assays, and after screening >40,000 EMS-mutagenized chromosomes, we identified multiple alleles of only three genes (dSarm, axundead, and highwire) that provided protection similar to *dSarm* null mutants. This implies that even in the simplified genome of *Drosophila*, there are only a handful of genes that can exert this remarkably protective effect on axons after axotomy.

In this study we turned to dNmnat depletion, a well-known activator of dSarm/Sarm1. In contrast to axotomy, this approach leaves the entire neuron intact, dNmnat loss activates dSarm signaling, and this the entire cell undergoes neurodegeneration. It is possible that this approach might better model neurodegenerative diseases where axons are not severed, but dNmnat depletion often occurs and ultimately activates dSarm/Sarm1 signaling. In principle, this approach enables genetic interrogation of pro-degenerative signaling in both the soma and axon, and while we did not identify mutants that affected either compartment selectively, such molecules might exist, and this method holds promise to identify them. Accumulating genetic evidence supports the notion that the genetic pathways that drive neurodegeneration after axotomy versus dNmnat depletion could be distinct. For instance, mutants like *dWnk*^15^ can strongly block neurodegeneration activated by dNmnat^RNAi^, but not axotomy. Indeed, this was true of nearly all the suppressors we identified of neurodegeneration after dNmnat depletion. Despite this fact, an exciting possibility is that by generating double- or triple-mutant combinations of these lines, we might find combination exhibiting neuroprotection in axotomy similar to *dSarm* nulls. It will also be exciting to also explore the potential neuroprotective effect of these mutants in *Drosophila* models of neurodegeneration.

### CG4098/Nudix9 and NAD^+^ metabolism in axon protection

One of our strongest hits, *CG4098*^*KO*^, encodes the *Drosophila* homolog of human NUDT9, a nudix hydrolase that hydrolyzes ADPR and cADPR^45^. Loss of *CG4098* protected axons from dNmnat depletin and causes alterations in metabolite profiles, including accumulations of ADPR and cADPR as well as changes in NAD^+^ precursors such as NA and NaR (Fig. 3B). Of note, NAD^+^ levels remain unchanged in all conditions – suggesting that protection is not simply explained by preservation of NAD^+^ itself but may instead reflect changes in metabolite flux or signaling intermediates. Changes in NAD^+^ signaling intermediates may for example make neurons more resistant to drops in Nmnat.

It was surprising to identify a genetic manipulation that increases cADPR also protects axons from dNmnat depletion. In mammals, manipulation of cADPR alone has not been sufficient to alter the kinetics of axon degeneration in mammals after axotomy, but it does serve to some extent as a biomarker for SARM1 activity^46^. Moreover, genetic or pharmacological inhibitors of cADPR production? prevent axonal degeneration from paclitaxel toxicity – a common chemotherapeutic drug^47^. Although it is a byproduct of NAD^+^ hydrolysis by dSarm/Sarm1, how well cADPR serves as a biomarker for dSarm/Sarm1 activity *in vivo* is not clear. Because we measured metabolites in complex tissues, there could be cell-specific changes in cADPR across cells, so neuroprotection might not correlate its levels. In addition, precisely how *CG4098* loss alters metabolism in wing sensory neurons (versus the visual system) remains unknown. It is also possible that neuronal metabolic profiles could change drastically when dNmnat levels are depleted. Nevertheless, our findings align with emerging evidence that NAD^+^ metabolism exerts multifaceted effects on neuronal health and that different nodes of this pathway can differentially influence degeneration.

### Abrupt regulates dNmnat levels to preserve axons

Our screen also identified loss-of-function alleles of *abrupt*, a BTB-zinc finger transcription factor previously implicated in neuronal differentiation^37,38^. We find that *ab* mutants robustly suppress degeneration following dNmnat depletion, and this protection correlates with increased dNmnat protein levels (Fig. 4A, C). The presence of an Abrupt binding motif in the 5’ UTR of dNmnat suggests a direct regulatory relationship, and we speculate that Abrupt acts as a negative regulator of dNmnat expression, with its loss boosting dNmnat levels thereby enhancing axonal survival. This mechanism would parallel the well-established protection conferred by *Wld*^*S*^, a fusion protein that results in sustained high Nmnat levels in axons and thereby suppression of neurodegeneration. Thus, both genetic loss of Abrupt and Wld^S^ expression converge on the same principle: enhanced availability of Nmnat stabilizes axons and delays degeneration.

## Methods

### Fly Husbandry

Flies were maintained on standard cornmeal-agar food at 25^°^C under a 12:12 h light/dark cycle. For experiments requiring enhanced Gal4/UAS activity, flies were reared at 29°C. Unless otherwise noted, experiments were performed in the w^1118^ background.

### Generation of neuronal MARCM Clones

Neuronal clones were generated using Mosaic Analysis with Repressible Cell Marker (MARCM) as previously described^14^. Briefly, *ase-FLP* was used to induced mitotic recombination at FRT sites in combination with neuron specific Gal4 drivers. We employed pan neuronal (*elav-Gal4, Appl-Gal4*), glutamatergic (*OK371-Gal4*), and cholinergic (*ChAT-Gal4*) drivers depending on which chromosomal arm we induce clones. MARCM clones expressed *UAS-dNmnat RNAi* and *UAS-mCD8::GFP*, enabling simultaneous knockdown of dNmnat and visualization of affected neurons.

### Induction of Axon Degeneration

To trigger degeneration, *UAS-dNmnat RNAi* constructs were expressed in MARCM clones. Degeneration was scored 10 days post-eclosion (DPE), when RNAi induction produced robust and reproducible axonal loss.

### Confocal imaging and Processing

Imaging of the adult wing nerve was performed as previously described^14^ with minor adjustments. Briefly, 10-day-old flies were anesthetized with CO_2_ and wings were removed using spring scissors. Wings were mounted in Halocarbon Oil 27, under #1.5 coverslips and imaged within 20 min of mounting at room temperature. Z-stack images spanning the L1 longitudinal vein were acquired on a Zeiss Axio Examiner with a Yokogawa spinning disk and Hamamatsu camera using a 63×/1.4 NA oil-immersion objective with GaAsP detectors (488-nm excitation for GFP). Laser power, detector gain, pinhole (1 AU), pixel size, and z-step (0.3–0.5 µm) were kept constant across samples within an experiment, and control and experimental wings were imaged in the same session. Image acquisition used Zeiss ZEN Black; processing was performed in Fiji/ImageJ and included maximum-intensity projections; minimal processing which included rotation and cropping only. No nonlinear adjustments were applied; any linear brightness/contrast adjustments were applied equally to all images in a set.

### EMS mutagenesis and F1 Screen

Forward genetic screening was performed using ethyl methanesulfonate (EMS) mutagenesis. P_0_ males were mutagenized with 25 mM EMS in 1% sucrose and crossed to virgin tester females carrying the appropriate MARCM insertions. F1 progeny generated homozygous mutant MARCM clones in the wing while maintaining heterozygosity in non-clonal cells, permitting assessment of clone-specific phenotypes. Candidate suppressor lines were maintained through balancer chromosomes. In total, 9393 mutagenized chromosomes across all major chromosomal arms were screened.

### Whole-Genome Sequencing

Genomic DNA was extracted from pooled flies of candidate lines using Thermo Fisher DNA extraction kit. Illumina short-read sequencing was performed, and variants were called using GATK and compared against the reference *Drosophila melanogaster* genome (BDGP6). Candidate mutations were prioritized based on predicted coding consequences.

### CRISPR/Cas9-Mediated Gene Editing

We generated a 358 bp deletion and adenine insertion in *CG4098*. sgRNAs targeting exon 2 were cloned into pU6BbsI and injected into *y1M{vas-Cas9}ZH-2A* embryos. Deletions were verified by PCR and sanger sequencing provided by GENEWIZ.

### Metabolite Profiling

Heads from 3–5 day old adult flies (50 per sample) were flash frozen in liquid nitrogen and extracted with ice-cold methanol containing stable isotope–labeled internal standards. Extracts were centrifuged, dried under vacuum, and reconstituted in ddH_2_O for LC–MS/MS analysis. Metabolites were separated on a Scherzo SMC18 column and quantified on an Agilent 6495 triple quadrupole mass spectrometer using dynamic multiple reaction monitoring in positive electrospray ionization mode. Concentrations of NAD^+^, NMN, ADPR, cADPR, NA, NaR, and related intermediates were determined from calibration curves, normalized to internal standards, and corrected for total protein.

### ELISA assay

dNmnat protein levels were quantified using a GFP-based ELISA adapted from established protocols^42^. Adult fly heads were homogenized in 1XPBS with 0.1% Tween-20 (PBST). High-binding 96-well plates were coated with anti-GFP antibody, blocked with 2% milk in PBST, incubated with lysates, and detected using HRP-conjugated secondary antibodies with TMB substrate. Absorbance at 450 nm was measured, and dNmnat-GFP concentrations were interpolated from the standard curve and normalized to total protein input.

### Statistics

Statistical analyses were conducted in GraphPad V10. For multiple comparisons, one-way ANOVA was used. P≤0.05 is *, P≤01 is **, P≤ 0.001 is ***, P≤0.0001 is ****.

