## Supplemental Figures for "A dNmnat-sensitized *in vivo* platform for unbiased discovery of regulators of neurodegeneration"

Supplementary figures:

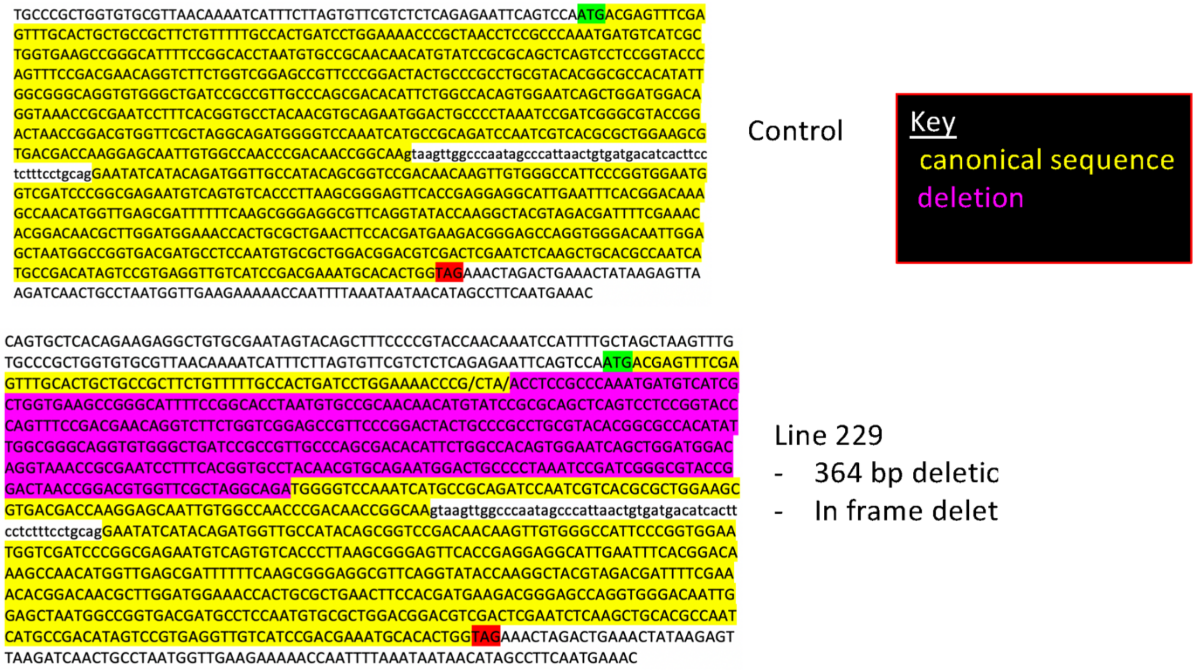


**Supplementary Figure 1. Sequence characterization of CG4098 mutant allele.** Comparison of wild-type (control) and EMS mutant line 229 sequences. The mutant allele contains a 364 bp in-frame deletion corresponding to 118 amino acids. Canonical coding sequence is highlighted in yellow, and the deleted region is marked in magenta. This deletion preserves the reading frame but removes a large internal segment of the encoded protein, consistent with a functional loss-of-function allele.

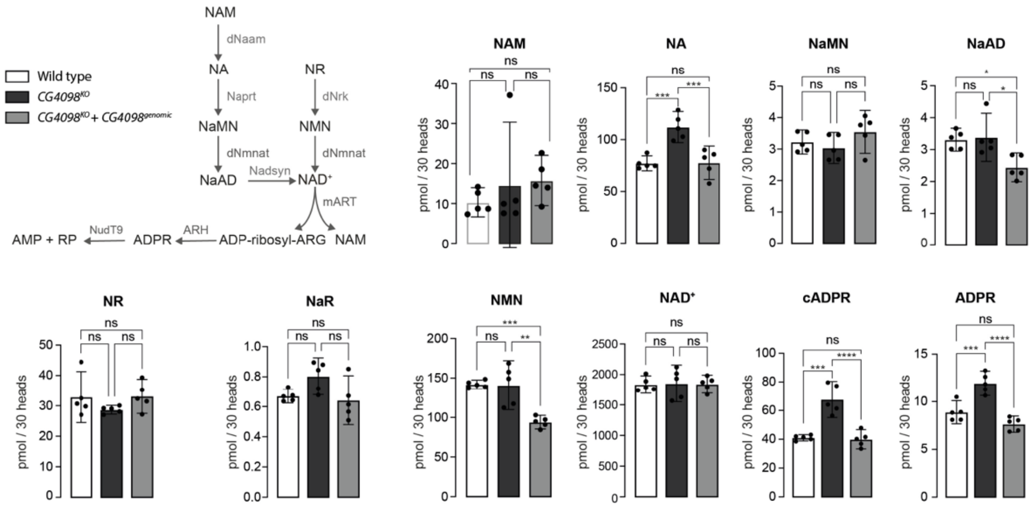

**Supplementary Figure 2. Expanded metabolite profiling of CG4098/NUDT9 mutants.** (A) Schematic of the $NAD^+$ biosynthetic and degradation pathways highlighting all metabolites quantified.

(B) LC–MS/MS analysis of metabolites from adult heads comparing wild type, $CG4098^{KO}$, and *CG4098*$^{KO}$ + *CG4098*$^{genomic\ rescue}$ flies. In addition to the metabolites shown in the main figure (NMN, NA, $NAD^+$, cADPR, ADPR), expanded profiling revealed no significant changes in nicotinamide (NAM), nicotinic acid mononucleotide (NaMN), or nicotinamide riboside (NR). Nicotinic acid adenine dinucleotide (NaAD) was modestly reduced in rescue animals compared to nudt9^184 mutants. As in the main figure, NMN, NA, cADPR, and ADPR levels were significantly elevated in nudt9^184 mutants and restored to wild-type levels in genomic rescue flies, while total $NAD^+$ levels were unchanged. Data represent mean ± s.e.m.; *p < 0.05, **p < 0.01, ***p < 0.001, ****p < 0.0001; ns, not significant (one-way ANOVA with multiple comparisons).
